# Multifunction control and evaluation of a 3D printed hand prosthesis with the Myo armband by hand amputees

**DOI:** 10.1101/445460

**Authors:** M. Cognolato, M. Atzori, C. Marchesin, S. Marangon, D. Faccio, C. Tiengo, F. Bassetto, R. Gassert, N. Petrone, H. Müller

## Abstract

Upper limb amputations are highly impairing injuries that can substantially limit the quality of life of a person. The most advanced dexterous prosthetic hands have remarkable mechanical features. However, in most cases, the control systems are a simple extension of basic control protocols, making the use of the prosthesis not intuitive and sometimes complex. Furthermore, the cost of dexterous prosthetic hands is often prohibitive, especially for the pediatric population and developing countries. 3D printed hand prostheses can represent an opportunity for the future. Open 3D models are increasingly being released, even for dexterous prostheses that are capable of moving each finger individually and actively rotating the thumb. However, the usage and test of such devices by hand amputees (using electromyography and classification methods) is not well explored. The aim of this article is to investigate the usage of a cost-effective system composed of a 3D printed hand prosthesis and a low-cost myoelectric armband. Two subjects with transradial amputation were asked to wear a custom-made socket supporting the HANDi Hand and the Thalmic Labs Myo armband. Afterwards, the subjects were asked to control and use the prosthetic hand to grasp several objects by attempting to perform a set of different hand gestures. Both the HANDi Hand and the Myo armband performed well during the test, which is encouraging considering that the HANDi Hand was developed as a research platform. The results are promising and show the feasibility of the multifunction control of dexterous 3D printed hand prostheses based on low-cost setups. Factors as the level of the amputation, neuromuscular fatigue and mechanical limitations of the 3D printed hand prosthesis can influence the performance of the setup. Practical aspects such as usability and robustness will need to be addressed for successful application in daily life. A video of the tests can be found at the following link: https://youtu.be/iPSCAbd17Qw

## I. INTRODUCTION

Upper limb amputations are highly impairing injuries, in particular if bilateral. Dexterous, naturally controlled robotic hand prostheses can substantially improve the quality of life of subjects with hand amputation. However, the control systems usually implemented into commercial devices are still limited and scientific research advancements are still poorly translated in commercial products [1], [2]. Low-cost resources, such as additive manufacturing and affordable surface electromyography (sEMG) sensors can foster a faster evolution of the field. Despite the many advancements, multifunction user tests of a 3D printed robotic hand prosthesis by hand amputees using low-cost devices have to the best of our knowledge not been presented, so far. In this paper, two subjects with transradial amputation use a dexterous 3D printed robotic hand prosthesis in multiple tasks selected from the activities of daily living. This indicates the feasibility of cost-effective systems composed of a 3D printed hand prosthesis and a myoelectric control system based on gesture recognition.

Poly-articulated dexterous myoelectric hand prostheses are now commercially available. Despite the remarkable mechanical capabilities of such devices, their control is still limited and often obtained as an extension of standard control protocols, making their use unintuitive and sometimes difficult [1], [3], [4], [2]. Several new control strategies were investigated in scientific research [2]. Invasive methods such as targeted muscle reinnervation (TMR) [5], [6], intramuscular electrodes [7], cortical implants [8] and peripheral nerve interfaces [9] have been developed and studied [1], [3], [2], [10], [4]. Machine learning techniques applied to myoelectric signals recorded from several electrodes placed on the forearm are arguably the most widely used non-invasive approaches investigated in the scientific research [11], [4]. Although some of these achievements can now be candidates for market translation, the implementation of scientific research advances into commercial products is still limited [1], [2]. Furthermore, advanced dexterous hand prostheses are usually very expensive (several tens of thousands of dollars [12], [13]) and healthcare systems or insurance reimbursement policies do not systematically cover the costs completely. Hence, the economic aspect can sometimes be crucial, in particular in settings such as pediatric populations or in developing countries, where hand amputations are more frequent as well.

In recent years, inexpensive devices and technologies such as open-source electronic prototyping platforms and 3D printing techniques have been successfully applied in the field of prosthetics. Thanks also to communities such as e-NABLE^1^, Limbitless Solutions^2^, and Open Hand Project^3^, several models of 3D printable hand prostheses are now publicly available. Open-source and inexpensive electronic platforms such as Arduino^4^ can be used to implement low-cost myoelectric control systems.

3D printed hand prostheses represent an opportunity and are available but only rarely tested in functional settings with more intuitive control systems. Commercial prosthetic hands are commonly very expensive and limitations such as control difficulties and comfort problems are common causes of prosthesis rejection [14], [15], [16]. 3D printed prostheses currently represent an affordable resource that can easily and quickly be adapted to the subject’s needs as well as to scientific advancements. For these reasons, several groups and communities release open-source designs, with various characteristics and complexity, including models targeting essentially research needs [16].

The high cost of high-end technical equipment and sensors can create a barrier that is not easy to overcome for research groups that are located in developing countries or that have limited funding. This was the case of robotic hand prostheses control, as well. A few years ago, a low-cost gesture recognition armband called Myo was released by Thalmic Labs (Ontario, Canada)^5^, costing approximately 200 dollars instead of other control systems that costs in the range of $10,000. The Myo armband was successfully applied in hand gesture recognition tasks in intact and hand amputated subjects [17], [18], [19], [20]. The performance of the Myo for gesture classification tasks was compared with 5 other commonly used acquisition setups on a standardized data acquisition and analysis protocol, showing that the classification accuracy is comparable to expensive setups in a two-armband configuration and moderately lower in the standard configuration [18].

Although low-cost devices and techniques are making prosthetic hands more accessible, improvements are still needed. First, 3D printed hand prostheses often have limited mechanical features with respect to commercial devices, restricting their applicability for daily use. Second, the use of a prosthetic hand in a real-life scenario can be substantially influenced by factors such as clinical parameters of the amputee [21] and comfort questions. Furthermore, models of 3D printable hand prostheses are often released “as-is” and the user is responsible for evaluating potential risks e.g. mechanical failures.

To the authors’ knowledge, this paper presents one of the first multifunction control tests of a dexterous 3D printed prosthetic hand in activities of daily living performed by subjects with transradial amputation using a control system based on gesture recognition. The results are encouraging and highlight the advantages and disadvantages of the current state of the art, suggesting the feasibility of cost-effective setups composed of a 3D printed hand prosthesis and affordable myoelectric control systems based on gesture recognition.

## II. METHODS

In this section the characteristics of the subjects, the acquisition setup and protocol are presented along with the real-time control system and the evaluation tasks. The steps described allow subjects with transradial amputation to control the 3D printed hand prosthesis with multiple functions and to test the entire system in conditions simulating a real life environment.

### A. Subjects

Two male transradial amputees were recruited for this study, both with traumatic injury as the cause of the amputation (Fig. 1). Subject 1 lost his dominant hand (right) in 1999, while subject 2 had the amputation of his non-dominant hand (left) in 2000. The level of the amputation is very different for the two subjects: subject 1 has a wrist disarticulation while subject 2 has a high-level transradial amputation. In order to allow the comparison between different subjects, the length of the residual limb was normalized with respect to the length of the intact (contralateral) forearm and quantized in 5 segments of 20% width. The mean value of the correspondent segment is reported. Both subjects use a one degree of freedom (DoF) hand prosthesis in their daily life. Subject 1 uses a myoelectric hand prosthesis, while subject 2 uses a body-powered prosthetic hand. Detailed information about the subjects are reported in Table I.

**Fig. 1:**
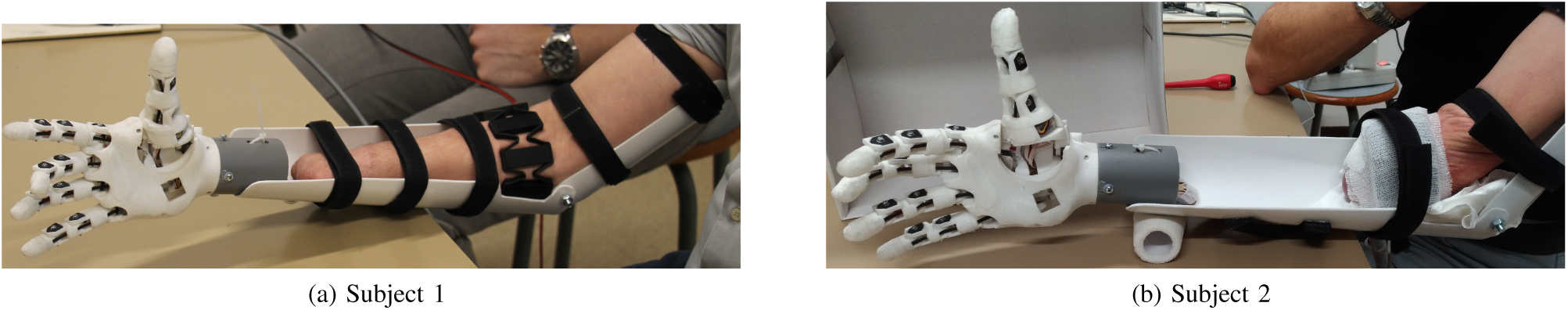
Acquisition setups for Subject 1 and Subject 2.

**TABLE I:**
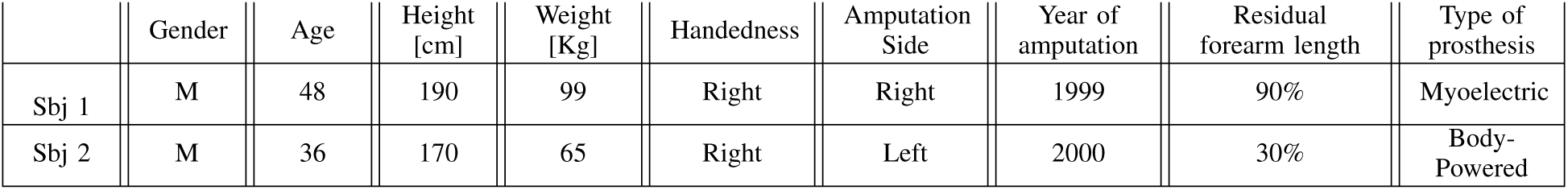
Characteristics of the subjects with hand amputation participating in this experiment.

### B. Acquisition setup

The acquisition setup is composed of:

- HANDi Hand 3D printed hand prosthesis [22]
- Myo, Thalmic Labs^5^
- Arduino Leonardo board^4^
- MyoDuino software^6^
- Custom-made socket
- Acquisition laptop

The 3D printed hand used in the tests is the Humanoid, Anthropometric, Naturally Dexterous Intelligent (HANDi) Hand, developed and released by the Bionic Limbs for Improved Natural Control (BLINC) Lab at the University of Alberta [22]. It is a low-cost, poly-articulated, sensorized 3D printed prosthetic hand designed to provide a cost-effective platform for machine learning research in hand prostheses myoelectric control. It is actuated by six servomotors: five controlling the flexion-extension of each finger and one actuating the thumb abduction/adduction. The angle at each finger joint is measured with potentiometers and the force at each fingertip is provided by force sensitive resistor (FSR) sensors. The mechanism used to flex and extend the fingers is based on cable ties and pulleys. Finger flexion is obtained by rolling the cable tie around a pulley attached to a servomotor, while the finger extension is obtained by simply unrolling it, using its residual flexural stiffness. Care was taken to choose the appropriate cable tie stiffness in order to maintain a satisfactory grip efficiency. The HANDi Hand weighs ~250 g and it has a maximum load of 500 g. The hand used in this experiment was printed in Nylon using selective laser sintering (SLS) as the 3D printing technique. Some modifications were made to improve the robustness of the HANDi Hand and the *D-shaped inserts* (mechanical parts transmitting the motion of the finger to the potentiometers) were re-designed to solve a problem on the measurement of the joint angles on our hand, probably due to backlash.

The Myo^5^ is a wearable gesture recognition armband composed of sEMG electrodes and an inertial measurement unit (IMU). The IMU is composed of three-axis accelerometer, gyroscope, and magnetometer. An elastic framework containing eight medical grade stainless steel dry sEMG electrodes form the armband, which can fit a forearm circumference ranging from 19 to 34 centimeters. The Myo communicates wirelessly via Bluetooth, it is lightweight (93 g) and it has a built-in real-time hand gesture classifier that allows it to be used as gesture and motion control. Five pre-set hand gestures are recognized by the Myo, namely, *fist, wave in* (wrist flexion), *wave out* (wrist extension), *fingers spread*, and *double tap*. The first four gestures are purely EMG-based whereas orientation and rotation data from the IMU are used to identify when the *double tap* gesture is triggered.

The HANDi Hand is controlled via the open-source electronic prototyping platform Arduino Leonardo^4^. Each servomotor is connected to a pulse width modulation (PWM) port of the board and controlled using the Arduino *Servo* library. The *MyoDuino*^6^ software was used to read and send the hand gesture identified by the Myo to the Arduino board in real-time. The connection between the Myo and Arduino was performed via a laptop, connected to the Arduino via USB cable and to the Myo via Bluetooth.

A light-weight custom-made socket designed to be used with various sizes of the remaining forearm was used in this experiment. The custom-made socket is composed of three parts: a forearm support, an arm support and a connector for the HANDi Hand. The forearm and arm supports were connected at the elbow height, allowing the wearers to flex and extend the elbow. Both forearm and arm supports were fixed to the patients limb using velcro straps and the HANDi Hand was attached to the socket by mean of a connector placed at the end of the forearm support (Fig. 1).

### C. Acquisition protocol

The acquisition protocol consisted of grasping various objects several times with the 3D printed prosthetic hand, controlling the 3 hand grasps. The hand grasps were chosen among the most frequently used grasps in the activity of daily living (ADL) tasks [23], [24]. They are: *medium wrap, power sphere,* and *lateral pinch*. Each grasp was performed on a set of three objects (Fig. 2).

**Fig. 2:**
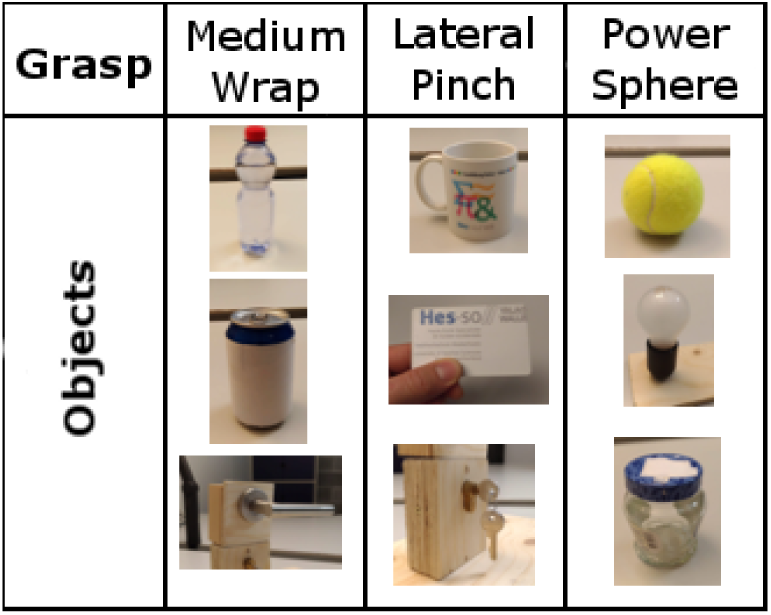
Objects used in the data acquisition protocol to perform the activities. The subject had to grasp the three objects four times with the correct grasp.

At the beginning of the experiment, the subjects received a detailed description of the experiment and were asked to sign an informed consent form. The sEMG acquisition protocol was developed according to the principles expressed in the Declaration of Helsinki and approved by the Ethics Commission of the Province of Padova (Italy). Afterwards, the subjects were asked to wear the Myo armband and to complete the algorithm calibration for the *fist, wave in, wave out* and *fingers spread* gestures. The subjects were then asked to freely try the calibrated grasps while receiving a visual feedback indicating which hand gesture was being recognized by the Myo. Once the subject felt ready, he was asked to wear the custom-made socket with the HANDi Hand.

Since the position of the Myo armband may have changed while donning the socket, the calibration was repeated. The subjects were asked to comfortably sit in front of a desk, and, for each grasp type, they were requested to use the HANDi Hand to grasp four times a set of three objects. Each object was positioned in front of the subject individually. For each grasp and object combination, one examiner asked the subjects to attempt performing the requested grasp while another examiner annotated the time needed to complete the action and the number of misclassifications. Each trial ended when each object was correctly grasped for four times. The tests were performed at the Sports & Rehabilitation Laboratory of the Department of Industrial Engineering, University of Padova.

### D. Real-time control

The real-time control of the HANDi Hand was based on the hand gesture classification performed by the Myo armband classifier. Thus, each gesture of the Myo triggers a specific movement of the HANDi Hand in real-time. The real-time control system was designed to be easy-to-use and as little tiring as possible for the subjects. When a hand gesture was identified by the Myo classifier, the HANDi Hand changed the grasp and no more muscular activity was required. In order to change a grasp, the HANDi Hand had first to be opened performing the *fingers spread* grasp.

For each grasp, the shape of the HANDi Hand was pre-set as follows:

- *Fist* → *medium wrap:* the thumb is first adducted, then all the fingers are flexed.
- *Fingers spread* → *opening:* the thumb is completely abducted and all the fingers fully extended.
- *Wave in* → *power sphere:* all the fingers are flexed and the thumb adducted at the same time.
- *Wave out* → *lateral pinch:* first index, medium, ring and little fingers are flexed and, after a delay of 2 seconds, the thumb is adducted and flexed.

## III. RESULTS

The results show the effectiveness of the approach presented in this work: a cost-effective setup consisting of a 3D printed dexterous prosthetic hand and the Myo armband can be used by subjects with transradial amputation to grasp several objects (Fig. 3).

**Fig. 3:**
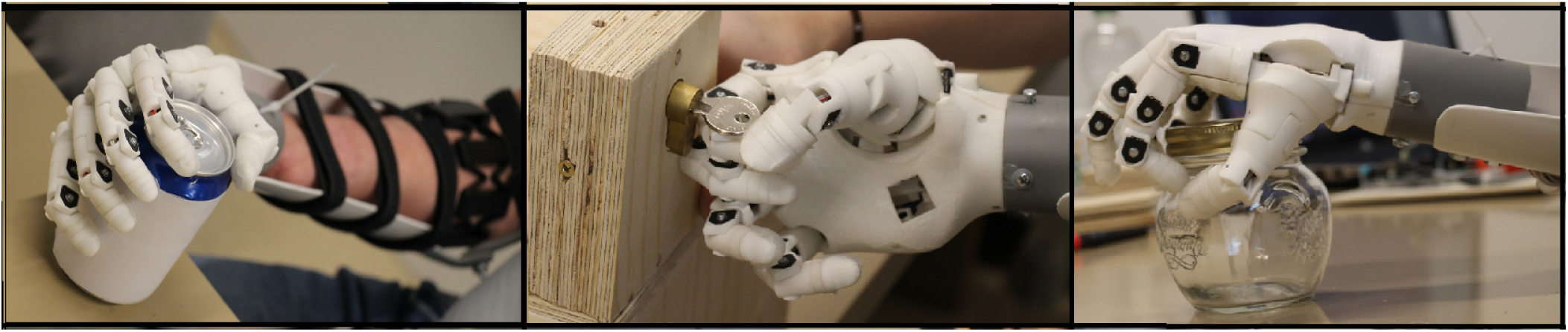
The selection of hand grasps performed by the subjects. From left to right the *medium wrap, lateral pinch* and *power sphere* grasps.

The myocontrol system allows the subjects to choose the grasp to be performed among various grasps, the number of which can be adapted to the user characteristics. Both subjects were able to successfully control the prosthetic hand using the proposed control system and reported the control system to be comfortable in most of the considered grasps.

The HANDi Hand was designed targeting scientific research applications rather than everyday use [22]. Nevertheless, the mechanical characteristics of this hand make it potentially suitable to perform essential everyday tasks such as lifting a 0.5l water-bottle as well as pressing a door handle to open it. The HANDi Hand performed well on each task and the subjects highlighted several advantages with respect to their prosthesis, including the high dexterity and the light weight (which is lower than the weight of standard myoelectric hand prostheses). On the other hand, some technical problems were experienced, suggesting that improvements are still important for everyday life use. The servomotor gearbox had two failures during the experiments, highlighting a limitation of small actuators. Furthermore, three cable ties broke while testing the equipment as well as during the experiment.

The custom-made socket was designed to be low-cost, lightweight, simple to build, useful in scientific research applications and usable by transradial amputees with various level of amputation as well as intact subjects. Both subjects reported that the custom-made socket was easy to wear and effective for the experiments. On the other hand, the leverage applied by the socket on the residual limb can limit the comfort for subjects with a high-level amputation. In particular, subject 1 was able to sustain the weight whereas the same was not possible for subject 2 with this arrangement. In addition, no artifacts in the Myo data were noticed while the armband was worn inside the custom-made socket.

The Myo allows performing myoelectric control using a classification approach, as it is often done in scientific research. The sEMG signals are recorded with several electrodes around the residual limb and the hand gesture is classified based on these data. The myocontrol system proposed in this work allows the identification of four hand gestures and the control of three grasps. The real-time control was designed to perform the grasps that are frequently used in ADL with the minimum cognitive and physical load for the subject. The shape of the HANDi Hand for each grasp was pre-set and triggered as soon as a hand gesture was identified by the Myo classifier. Thanks to this approach the subject was free to focus on performing the correct hand gesture, with no need of continuous muscular contraction. Furthermore, the constraint requiring the hand prosthesis to be opened before switching to another type of grasp assured a good stability, preventing undesirable hand behaviour due to involuntary muscular activation. Performing the hand gestures was reported as comfortable by subject 1 whereas subject 2 highlighted a few difficulties while attempting to perform the movement. Nevertheless, both subjects were able to control the prosthesis after less than an hour of training, while most commercial systems require several days (or, more often, weeks) of training.

Subject 1 was able to control and use the 3D printed hand in all the requested grasps. On average, the time needed to grasp each object with the *medium wrap* and *power sphere* grasps was 2 seconds. A longer time was needed to perform the *lateral pinch* grasp, due also to the time with which the shaping of the grasp was programmed. Indeed, the *lateral pinch* grasp had an intrinsic delay of 2 seconds, in order to allow the subject to properly exploit the side opposition required by the grasp. The *medium wrap* gesture trial was completed by subject 1 in 22 attempts, with a misclassification rate of 45.45%. For the same subject, the *power sphere* grasp trial was misclassified only once (13 attempts with a misclassification rate of 7.69%) whereas eighteen tries were needed to complete the *lateral pinch* gesture task, with a misclassification rate of 33.33%. Two main factors were identified as the cause of misclassification: changing in limb posture and neuromuscular fatigue.

Subject 2 has a high-level amputation (Fig. 1b), with ~ 30% of residual limb length. This factor limits the capabilities of the subject to the point that he cannot use a standard myoelectric prosthesis. This also limited the custom-made socket fitting as well as the Myo armband position. Due to the small remaining forearm, using the socket and the HANDi Hand was not comfortable for the subject. Therefore, he was asked to control the hand while the socket was leaning on the table and the objects were placed in front of the HANDi Hand in order to be grasped. For subject 2, the hand gesture recognition system had a few limitations. The *fingers spread* gesture was often misclassified as *wave in*, while the *wave out* gesture was reported as difficult and tiring to perform. Hence, the real-time control of the hand was modified accordingly to the subject’s characteristics: both *fingers spread* and *wave in* were linked to the opening of the HANDi Hand and the *wave out* gesture was excluded from the trials. This limited the number of hand gestures to two: *fist* and *fingers spread & wave in,* allowing the subject to perform the *medium wrap* grasp and to open the prosthetic hand. On the other hand, this approach increased control robustness, reduced mental and physical stress and made the subject more confident. The time needed for subject 2 to grasp an object is similar to the ones obtained by subject 1 and he performed all the 12 grasps without any classification error.

## IV. DISCUSSION

The results suggest the effectiveness of dexterous robotic prosthetic hands based on cost-effective setups composed of a 3D printed hand prosthesis and a myoelectric gesture classifier (about 100 times less expensive than advanced commercial products).

An increasing number of 3D printed hand prosthesis models are now publicly available. The load and mechanical robustness of commercial prostheses are often higher than the ones provided by 3D printed devices. Nevertheless, 3D printed hand prostheses can restore basic but essential hand functionalities, improving the quality of life of subjects with amputation. Despite being designed targeting scientific research applications, the mechanical features of the HANDi Hand allow it to perform several daily life activities. Furthermore, the HANDi Hand can provide contextual information about both the prosthesis and the surrounding environment. These additional sources of information can be used in multimodal machine-learning-based myoelectric control systems as well as to provide sensory feedback to the user, with great potential also for a real use—for example, the control algorithm can retrieve information about the object to grasp from the in-palm camera and a mechanical pressure proportional to the force sensed at the fingertips can be provided as sensory feedback. On the other hand, the technical problems experienced during the experiments suggest that a few improvements are required to obtain a sufficient level of robustness and reliability for a sustained everyday life use.

The myoelectric control system was based on the Myo armband and its gesture classifier. As previously suggested in [17], this work shows that subjects with transradial amputation can use the Myo armband and its classifier to control a dexterous robotic hand prosthesis. The quality of the control was different between the two subjects, mainly due to clinical parameters such as remaining forearm percentage (as reported in [21]). The system can also be improved by applying customizable calibration and machine learning techniques to the raw sEMG data acquired from the Myo. This approach can give full control on a higher number of hand gestures, making the system fully customizable according to the characteristics of the subject.

The myocontrol system proposed in this study differs from the approach usually implemented in commercial devices. While in traditional control systems the grasp to be performed is usually pre-selected by the user, in the proposed approach the 3D printed prosthetic hand is controlled by hand gestures. For instance, in the proposed control system the opening and closing of the prosthesis are performed with the *fingers spread* and *fist* gestures. According to the test subjects, this approach makes the proposed control system more intuitive than the ones usually implemented in commercial devices, as suggested by the fact that both subjects were able to successfully control the prosthetic hand with a short training phase.

Subject 1 was able to perform all the four hand gestures and grasping the objects with an average grasping time of 2 seconds. The level of the amputation of subject 2 makes traditional myoelectric control a challenging task for him. The difficulties in finding two optimal sEMG electrode locations and the height of the amputation prevented him from using a commercial myoelectric prosthetic hand. Nevertheless, the control system based on the Myo classifier was able to correctly and robustly distinguish between two sets of hand gestures, allowing him to control the 3D printed hand prosthesis.

Subject 1 is currently using a one DoF non-dexterous myoelectric hand prosthesis whereas subject 2 a body-powered device. The participants identified the ability to perform various grasps and potentially move each finger independently as a valuable advantage over the prostheses they use in daily life.

## V. CONCLUSIONS

The results are promising and suggest the feasibility of dexterous robotic prosthetic hands based on affordable setups. A 3D printed prosthetic hand was successfully controlled by the classifier provided with the low-cost Myo armband and used for daily life activities by 2 subjects with hand amputation. The gesture recognition approach implemented in the control system used in this study allowed the automatic identification of the desired grasp quickly (average grasping time below 3 seconds for most repetitions). The system was customized following the characteristics and needs of the subjects, allowing them to control up to 3 hand grasps in addition to the gesture used to open the prosthetic hand.

Several improvements can enhance the capabilities of the proposed control system. The application of machine learning techniques to the raw sEMG data acquired from the Myo can expand the capabilities of the myocontrol. Furthermore, the additional information provided by the sensors embedded in the HANDi Hand can be used to improve the control strategy as well as to give sensory feedback to the subject.

Thanks to the increasing affordability of both the components and devices, this technology is now more accessible than in the past. The overall cost of the entire setup (excluding the acquisition laptop that can be easily replaced by other devices such as a smartphone or even removed) is approximately 800 euros. This fact highlights that the setup used in this experiment can be reproduced and successfully used even where the cost is a critical factor, such as in the pediatric population, in developing countries and in research groups with limited resources.

## ACKNOWLEDGMENT

The authors would like to thank the Orthomedica orthopedic center, the subjects participating in this study as well as the Swiss National Science Foundation for partially supporting this work via the Sinergia project # 410160837 MeganePro. Furthermore, the authors would like to thank D. J. A. Brenneis, M. R. Dawson and P. M. Pilarski for the development and release of the HANDi Hand.

## Footnotes

1 http://enablingthefuture.org/

2 https://limbitless-solutions.org/

3 http://www.openhandproject.org/

4 http://www.arduino.cc/

5 http://www.myo.com/

6 https://market.myo.com/app/54bd7403e4b00db53ad527a2/myoduinoin

## REFERENCES

[1] D. Farina and S. Amsüss, “Reflections on the present and future of upper limb prostheses,” Expert Review of Medical Devices, vol. 13, no. 4, pp. 321–324, 2016.

[2] C. Castellini, P. Artemiadis, M. Wininger, A. Ajoudani, M. Alimusaj, A. Bicchi, B. Caputo, W. Craelius, S. Dosen, K. Englehart, D. Farina, A. Gijsberts, S. B. Godfrey, L. Hargrove, M. Ison, T. Kuiken, M. Marković, P. M. Pilarski, R. Rupp, and E. Scheme, “Proceedings of the first workshop on peripheral machine interfaces: Going beyond traditional surface electromyography,” Frontiers in Neurorobotics, vol. 8, no. AUG, pp. 1–17, 2014.

[3] S. Amsuess, P. Goebel, B. Graimann, and D. Farina, “Extending mode switching to multiple degrees of freedom in hand prosthesis control is not efficient,” in 2014 36th Annual International Conference of the IEEE Engineering in Medicine and Biology Society. IeEe, aug 2014, pp. 658–661. [Online]. Available: http://ieeexplore.ieee.org/document/6943677/

[4] M. Atzori and H. Müller, “Control Capabilities of Myoelectric Robotic Prostheses by Hand Amputees: A Scientific Research and Market Overview.” Frontiers in systems neuroscience, vol. 9, p. 162, 2015.

[5] T. A. Kuiken, G. A. Dumanian, R. D. Lipschutz, L. A. Miller, and K. A. Stubblefield, “The use of targeted muscle reinnervation for improved myoelectric prosthesis control in a bilateral shoulder disarticulation amputee.” Prosthetics and orthotics international, vol. 28, no. 3, pp. 245–53, dec 2004. [Online]. Available: http://www.ncbi.nlm.nih.gov/pubmed/15658637

[6] T. A. Kuiken, G. Li, B. A. Lock, R. D. Lipschutz, L. A. Miller, K. A. Stubblefield, and K. Englehart, “Targeted Muscle Reinnervation for Real-Time Myoelectric Control of Multifunction Artificial Arms,” JAMA, vol. 301, no. 6, pp. 619–628, 2009.

[7] C. Cipriani, J. L. Segil, J. A. Birdwell, and R. F. Weir, “Dexterous control of a prosthetic hand using fine-wire intramuscular electrodes in targeted extrinsic muscles,” IEEE Transactions on Neural Systems and Rehabilitation Engineering, vol. 22, no. 4, pp. 828–836, 2014.

[8] M. A. Lebedev and M. A. L. Nicolelis, “Brain-machine interfaces: past, present and future,” pp. 536–546, 2006.

[9] P. M. Rossini, S. Micera, A. Benvenuto, J. Carpaneto, G. Cavallo, L. Citi, C. Cipriani, L. Denaro, V. Denaro, G. Di Pino, F. Ferreri, E. Guglielmelli, K. P. Hoffmann, S. Raspopovic, J. Rigosa, L. Rossini, M. Tombini, and P. Dario, “Double nerve intraneural interface implant on a human amputee for robotic hand control,” Clinical Neurophysiology, vol. 121, no. 5, pp. 777–783, 2010. [Online]. Available: http://dx.doi.org/10.1016/j.clinph.2010.01.001

[10] M. Ortiz-Catalan, R. Brånemark, B. Hakånsson, and J. Delbeke, “On the viability of implantable electrodes for the natural control of artificial limbs: Review and discussion,” BioMedical Engineering Online, vol. 11, 2012.

[11] B. Peerdeman, D. Boere, H. Witteveen, R. Huis in ‘tVeld, H. Hermens, S. Stramigioli, H. Rietman, P. Veltink, and S. Misra, “Myoelectric forearm prostheses: State of the art from a user-centered perspective,” J. Rehabil. Res. Dev., vol. 48, no. 6, p. 719, 2011.

[12] D. K. Blough, S. Hubbard, L. V. McFarland, D. G. Smith, J. M. Gambel, and G. E. Reiber, “Prosthetic cost projections for service-members with major limb loss from Vietnam and OIF/OEF.” Journal of rehabilitation research and development, vol. 47, no. 4, pp. 387402, 2010.

[13] D. Van Der Riet, R. Stopforth, G. Bright, and O. Diegel, “An overview and comparison of upper limb prosthetics,” IEEE AFRICON Conference, 2013.

[14] D. J. Atkins, D. C. Heard, and W. H. Donovan, “Epidemiologic Overview of Individuals with Upper-Limb Loss and Their Reported Research Priorities,” JPO Journal of Prosthetics and Orthotics, vol. 8, no. 1, pp. 2–11, 1996.

[15] E. Biddiss and T. Chau, “Upper limb prosthesis use and abandonment: a survey of the last 25 years.” Prosthet. Orthot. Int., vol. 31, no. 3, pp. 236–257, 2007.

[16] J. ten Kate, G. Smit, and P. Breedveld, “3D-printed upper limb prostheses: a review,” Disability and Rehabilitation: Assistive Technology, vol. 12, no. 3, pp. 300–314, apr 2017. [Online]. Available: https://www.tandfonline.com/doi/full/10.1080/17483107.2016.1253117

[17] M. Cognolato, M. Atzori, D. Faccio, C. Tiengo, F. Bassetto, R. Gassert, and H. Müller, “Hand Gesture Classification in Transradial Amputees Using the Myo Armband Classifier,” in 7th IEEE RAS/EMBS International Conference on Biomedical Robotics and Biomechatronics (BioRob). Enschede, The Netherlands, 26 - 29 August 2018: In press, 2018.

[18] S. Pizzolato, L. Tagliapietra, M. Cognolato, M. Reggiani, H. Müller, and M. Atzori, “Comparison of six electromyography acquisition setups on hand movement classification tasks,” PLoS ONE, vol. 12, no. 10, pp. 1–16, 2017.

[19] A. Atasoy, E. Kaya, E. Toptas, S. Kuchimov, E. Kaplanoglu, and M. Ozkan, “24 DOF EMG Controlled Hybrid Actuated Prosthetic Hand,” pp. 5059–5062, 2016.

[20] S. Masson, F. Fortuna, F. Moura, and D. Soriano, “Integrating Myo armband for the control of myoelectric upper limb prosthesis,” XX V Congresso Brasileiro de Engenharia Biomédica, no. October, 2016.

[21] M. Atzori, A. Gijsberts, C. Castellini, B. Caputo, A.-G. M. Hager, S. Elsig, G. Giatsidis, F. Bassetto, and H. Müller, “Effect of clinical parameters on the control of myoelectric robotic prosthetic hands,” Journal of Rehabilitation Research and Development, vol. 53, no. 3, pp. 345–358, 2016.

[22] D. J. A. Brenneis, M. R. Dawson, and P. M. Pilarski, “Development of the Handi Hand: an Inexpensive, Multi-Articulating, Sensorized Hand for Machine Learning Research in Myoelectric Control,” 2017.

[23] I. M. Bullock, J. Z. Zheng, S. De La Rosa, C. Guertler, and A. M. Dollar, “Grasp frequency and usage in daily household and machine shop tasks,” IEEE Transactions on Haptics, vol. 6, no. 3, pp. 296–308, 2013.

[24] J. Z. Zheng, S. De La Rosa, A. M. Dollar, and S. De La Rosa, “An Investigation of Grasp Type and Frequency in Daily Household and Machine Shop Tasks,” proceedings of the 2011 IEEE International Conference on Robotics and Automation (ICRA), pp. 4169–4175, 2011. [Online]. Available: citeulike-article-id:9457399

